# Protein complex prediction using Rosetta, AlphaFold, and mass spectrometry covalent labeling

**DOI:** 10.1101/2022.04.30.490108

**Authors:** Zachary C. Drake, Justin T. Seffernick, Steffen Lindert

## Abstract

Structural mass spectrometry offers several techniques for the characterization of protein structures. Covalent labeling (CL) in combination with mass spectrometry can be used as an analytical tool to study and determine structural properties of protein-protein complexes. Degrees of modification obtained from CL experiments for specific labeled residues can be compared between the unbound and bound states of complexes. This analysis can yield insights into structural features of these protein assemblies, specifically the proximity of specific residues to the protein-protein interface. However, this data is sparse and does not unambiguously elucidate protein structure. Thus, computational algorithms are needed to deduce structure from the CL data. In this work we present a novel hybrid method that combines models of protein complex subunits generated with AlphaFold with differential CL data via a CL-guided protein-protein docking in Rosetta. In a benchmark set, the RMSD (root-mean-square deviation) of the best-scoring models was below 3.6 Å for 5/5 complexes with inclusion of CL data, whereas the same quality was only achieved for 1/5 complexes without CL data. The average improvement in RMSD observed upon inclusion of CL data was 5.2 Å. This study suggests that our integrated approach can successfully use data obtained from CL experiments to distinguish between nativelike and non-nativelike models.

**Significance Statement:** Structural mass spectrometry can be a powerful and versatile approach to characterize the structure of protein complexes. Data obtained from covalent labeling mass spectrometry can provide insights into higher order protein structure (particularly with respect to residue interactions and solvent accessibility) but needs to be supplemented by computational techniques to elucidate accurate, atomic-detail structural information. Here, we present a method to combine bioanalytical data obtained from covalent labeling with models generated using AlphaFold to accurately predict protein-protein complexes in Rosetta. Differential covalent labeling data can be used to determine the proximity of residues to the binding interface of complexes which we utilized to analyze computational models and improve structure prediction algorithms.

## Introduction

Mass spectrometry (MS) is a versatile analytical approach which has become a vital tool in structural biology, capable of probing the structure and dynamics of protein assemblies.^1, 2^ Protein-protein complexes are central in many crucial biological and cellular processes,^3^ which makes their structural elucidation important. Currently, over 182,000 protein structures have been determined and archived in the Protein Data Bank (PDB), with around 114,000 of these being protein-protein complexes.^4^ These high-resolution protein structures can be obtained using techniques such as nuclear magnetic resonance (NMR)^5^, cryo-electron microscopy (cryo-EM),^6^ and most notably X-ray crystallography.^7^ However, structures at atomic resolution are not always obtainable due to limitations of the forementioned techniques in areas such as acceptable system size, required sample concentration, and excessive sample conformational heterogeneity.

Structural mass spectrometry is an alternative method which generally requires less time for sample preparation, can handle smaller sample sizes, is usable for a large range of protein sizes, and provides sparse biophysical data that can be used to gain insight into a variety of protein structural characteristics. Although MS experiments cannot comprehensively determine a high-resolution protein structure, insights into conformational states can be obtained, validation of existing models can be achieved, or the data can be supplemented with computational techniques for atomic-detail structure elucidation. A few common approaches in structural MS are chemical cross-linking,^8^ hydrogen-deuterium exchange (HDX),^9, 10^ surface-induced dissociation (SID),^11, 12^ ion mobility (IM),^13^ and covalent labeling of macromolecules (CL).^14, 15^ Chemical cross-linking involves using chemical reagents that form covalent bonds to link specific functional groups within or across protein molecules, providing distance restraints. HDX methods can be used to study protein structure and dynamics using exchange between protein backbone amide protons and deuterium atoms from solution, which is sensitive to local solvent accessibility and flexibility. SID methods involve the soft ionization of native protein complexes into the gas phase which are then collided with a rigid surface where they can break apart into monomers or other intact subcomplexes. This method can provide information regarding the stoichiometry, connectivity, and interface strength of complexes. IM can offer structural information regarding the shape and size of a protein complex by analyzing the travel of a protein through a bath gas, providing an averaged cross-sectional area of the system. Finally, covalent labeling probes protein structure by exposing solvent-accessible amino acid side chains with either specific or nonspecific reagents that covalently bind. Differences in reactivity to labeling agents can distinguish between exposed and buried residues, as well as residues located at the surface of interacting domains in the case of protein complexes.

Covalent labeling offers several advantages over other MS techniques. For example, the challenging low abundance of specific interpeptide cross-links and complicated tandem MS fragmentation of chemically cross-linked peptides are not an issue for covalent labeling techniques.^16^ Furthermore, due to the formation of stable, covalent bonds, the labeling of amino acids with labeling reagents are usually irreversible, unlike HDX where back-exchange frequently occurs, adding additional layers of complexity. Sparse structural data can be obtained from covalent labeling experiments with reagents such as hydroxyl radicals, carbenes, trifluoromethylations (CF_3_), diethylpyrocarbonate (DEPC), and sulfo-N-hydroxysuccinimide acetate (NHSA).^17^ These experiments provide metrics of modification that depend on the reactivity, solvent accessibility and potentially other structural features of the specific residues in solution. Structures of protein complexes can be further probed by comparing the degree of modification in the unbound and bound states, when possible. Interface residues are generally identified by examining large changes in modification rates between the unbound/bound state of a complex as solvent accessibility is likely to most dramatically change at the protein-protein interfaces. For example, a residue that gets labeled readily in the monomer, but not in the complex is likely part of the interface; although protein-protein binding could cause tertiary conformational changes, which might also result in changes in modification. The data obtained through a covalent labeling MS experiment can thus be used to probe higher orders structure of protein complexes.

As an alternative to experimental methods, modern computational methods have seen great success in accurately predicting and modeling protein tertiary structure.^18, 19^ The recent release of AlphaFold2^20^ (AF2, from DeepMind) has resulted in a revolution in the accuracy of computational protein modeling. AlphaFold^21^ is a neural network-based model that takes advantage of sequence coevolution data which has shown remarkable success and has outperformed other prediction methods during the 13^th^ and 14^th^ (with AF2) Critical Assessment of Techniques for Protein Structure Prediction (CASP),^22, 23^ a series of blind tests to gauge the current state of protein structure prediction. AlphaFold-Multimer^24^ was released in 2021 and uses the AF2 model but was trained to predict multimeric complexes from sequences of multiple chains. Similarly, traditional protein-protein docking algorithms are useful for analyzing and predicting models of complexes. In protein-protein docking, monomeric structures (which can be obtained in a variety of ways) are used as input and structures of the complex are predicted, with favorable orientations of the different subunits. Existing protein-protein docking algorithms which have been successful include ClusPro,^25^ HDOCK,^26^ ZDOCK,^27^ SwarmDock,^28^ HADDOCK,^29^ PIPER,^30^ and RosettaDock.^31^ RosettaDock is a part of the Rosetta^32^ molecular modeling software suite which contains a large variety of algorithms for computational modeling and analysis of protein structures.

Incorporation of sparse experimental data into algorithms predicting protein structure can further improve computational predictions.^33–35^ Information obtained from hydroxyl radical footprinting (HRF), HDX, and DEPC labeling experiments have been shown to improve tertiary structure prediction with Rosetta^36–43^ by using calculated solvent exposure metrics for models to select for experimentally accurate predictions. Similarly, protein shape and size information obtained through collisional cross-section data from IM experiments has also improved Rosetta structure prediction.^44^ A method iSPOT,^45^ which uses a combination of multiple biophysical methods (integration of shape information from small-angle X-ray scattering and protection factors probed by hydroxyl radicals), has been shown as a powerful approach for integrated modeling of multiprotein complexes. Isotope exchange using HDX-MS has been used to improve protein complex prediction by simulating complex isotope patterns and comparing to those obtained experimentally.^46^ Similarly, differential HDX data has been incorporated into protein-protein docking to study the human uracil-DNA-gycosylase (hUNG) and its protein inhibitor (UGI).^47^ The use of differential covalent labeling has yet to be implemented within the RosettaDock framework. Although AF2 has proven to be an excellent and revolutionary method of protein structure prediction, there remain limitations, particularly for protein complexes.^48^ Covalent labeling has the potential to help overcome some of these limitations and Rosetta is uniquely suited for the development of hybrid methods incorporating labeling data. Here, we use RosettaDock to assemble protein complex subunits that were generated using AF2 and employ covalent labeling data to improve protein complex structure prediction.

In this study, we developed the computational framework for using covalent labeling data in protein complex modeling in cases when state-of-the-art methods (both AlphaFold-Multimer and Rosetta) underperform. We propose a score term dependent on differential covalent labeling data obtained from HRF, DEPC, or NHSA experiments which when combined with the Rosetta score function readily selects computational models which agree with experimentally determined structures. We first observed a correlation between differential modification rates and inter-subunit residue distances within a protein complex based on our structural hypotheses. Next, we developed a protocol where AF2 was used to generate structures of the protein subunits which were used as input for docking simulations. In a benchmark of 5 complexes, inclusion of our score term predicted 5/5 structures with root-mean-square deviation (RMSD) less than 3.6 Å when compared to the native crystal structure, as opposed to 1/5 without CL data.

## Methods

### Protein Complex Benchmark Set

The three protein complexes used in the benchmark dataset were actin bound to gelsolin segment 1 (actin/gs1, heterodimer, PDB ID: 1YAG),^49^ β-2-microglobulin (homodimer, PDB ID: 2F8O),^50^ and insulin (hexamer of heterodimers, PDB ID: 4INS).^51^ Crystal structures were available for each for the purpose of benchmarking predicted models. Residue-resolved differential covalent labeling data were also obtained for each system in both the unbound and bound states.^49–51^ The labeling reagent used for the actin/gs1 and insulin complexes was hydroxyl radicals and for β-2-microglobulin, the labeling reagents were diethyl pyrocarbonate (DEPC) and sulfo-*N*-hydroxysuccinimide (NHSA). There were 41 labeled residues for both the unbound and bound states for actin/gs1, 20 for β-2-microglobulin, and 17 for insulin. For benchmarking purposes, interface residues were defined as any residue with a heavy atom within 10 Å of a heavy atom in another protein subunit. Although each labeled residue had a measure for the frequency of modification in both the unbound and bound states separately, we wanted to directly quantify the change in modification between these states, hypothesizing that residues with large changes would likely be part of the protein-protein interface. For each complex in the data set, the modification change between different states of the complex was computed from the data, as shown in Equation 1, using the degree of labeling for each complex where *M_unbound_* and *M_bound_* are the degree of modification (modification rates for actin/gs1 and insulin, extent of modification for β-2-microglobulin) of the unbound and bound states of the complex, respectively.

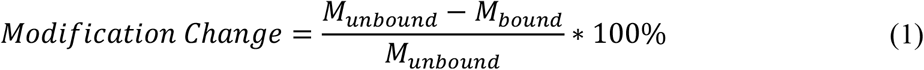

### Protein-Protein Docking

Docking simulations require input subunit structures which are used to predict the structure of complexes. In this work, we obtained input structures using two different methods. First, we used the bound crystal structures to perform a preliminary redocking study. Next, structures for each docking partner of actin/gs1 and β-2-microglobulin were generated using AlphaFold2 (AF2) for a more realistic prediction protocol.^20^ For insulin, the base subunit is a heterodimer, so AlphaFold-Multimer^24^ was used. The default settings for both AlphaFold methods were used along with the addition of all genetic databases (--db_preset=full_dbs flag). Since traditional docking consists of two docking partners, the insulin complex was broken up into three separate structures to model all unique interfaces, where AB_CD, ABCD_EFGH, ABCDEFGH_IJKL define what chains make up each docking partner, separated by an underscore (Figure S1). The docking protocol using RosettaDock was the same for each type of input structure. For each system, after prepacking, we generated sets of 10,000 docked models. The position and orientation of the second docking partner was randomized using the -randomize2 flag in the RosettaDock protocol to perturb each system.

### Complexes Generated using AlphaFold-Multimer

As a comparison to the docked models produced by RosettaDock, we also used AlphaFold-Multimer to predict full structures of each complex. To generate a more fair, blind prediction using AlphaFold-Multimer, restrictions were placed on which templates were used during model construction, as recommended by AlphaFold developers.^20^ We only included templates of structures with published dates prior to the date of the first published structure of each complex.

### Scoring Strategy

In this study, we proposed that differential covalent labeling data (comparing the unbound and bound states of a complex) could be used to indicate the proximity of a labeled residue to the binding interface of protein complexes and subsequently be used to assess model quality based on agreement with the experimental data. This was accomplished by comparing the modification change (Equation 1) of labeled residues and the distance from the interface in the crystal structures. The interface distance (Figure 1a) was defined as the shortest distance between a heavy atom of the target residue and a heavy atom from the binding partner. This comparison yielded an expected, inverse linear correlation between modification change and interface distance with the slope and intercept of the trendline being −2.07 and 46.27, respectively. This trendline was used for subsequent scoring.

**Figure 1:**
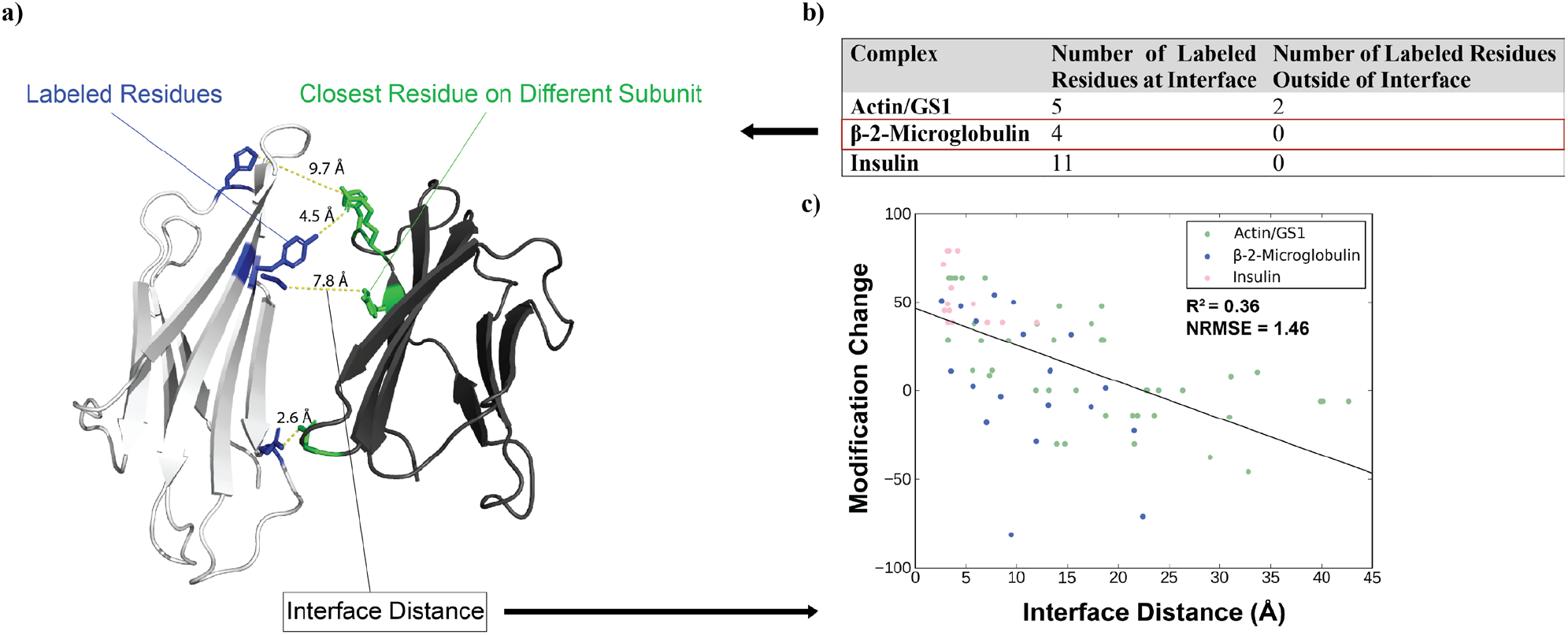
a) Visualization of the labeled residues with a modification change of at least 40% for β-2-microglobulin homodimer crystal structure. The labeled residues are colored in blue, their respective closest residue on a different subunit of the complex is colored in green, and the interface distance between them is shown above the dotted yellow line. b) Table indicating the number of residues with a modification change of at least 40%, within and outside the binding interfaces of a specified complex. c) Linear correlation between modification changes of labeled residues and the interface distances.

Therefore, to integrate the information regarding the modification change and interface distance into Rosetta to improve model scoring, a covalent labeling score term (*CL_Score_Term_*) was developed to assess the models generated with RosettaDock based on their agreement or disagreement with covalent labeling data. The covalent labeling score (*CL_Score_*), as defined in Equation 2, was the sum between the weighted *CL_Score_Term_* (described in the following paragraph) and the Rosetta Interface score (Isc).

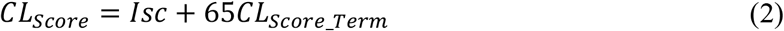

Isc was the energy of the binding interface of the docked complex calculated using the Rosetta REF2015 score function.^31^ *CL_Score_Term_*, defined in Equation 3, was a sum of per-residue penalties (*P_i_*) calculated using a sigmoidal penalty function. The penalty function scores labeled residues of a model based off deviations from the observed trendline of the native dataset (with large deviations from experimental results penalized).

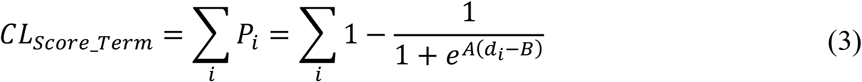

For each labeled residue of a given model, interface distance is calculated and used to predict modification change using the slope and intercept defined above. The difference (*d_i_*) between the experimental and predicted modification change was input into the penalty function. The A and B parameters defined the steepness and midpoint of the curve respectively, where A = 1.88 and B = 38.0. The summed penalties for all models are then normalized by dividing each score by the maximum score obtained for that particular system. Thus, the resulting *CL_score_term_* ranges from 0 to 1 where greater deviation from the trendline (indicating worse agreement with the experimental data) results in a larger penalty score from the score term.

### Analysis Metrics

The quality of models was assessed quantitively using alpha-carbon root-mean-squared deviation (RMSD), template modeling score (TM-score),^52^ and DockQ^53^ score with respect to the native crystal structure. For each model, the global RMSD values were calculated using PyMol.^54^ TM-score was used to analyze the topological similarity to the native structures. The TM-score ranges from 0.0 to 1.0 where a perfect match corresponds to a TM-score of 1.0. DockQ is a protein-protein docking quality measure which ranges between 0.0 and 1.0 with a perfect match being equal to 1.0.

## Results and Discussion

### Correlation of Differential Covalent Labeling and Residue Proximity to Binding Interface

We hypothesized that differential covalent labeling data could be used to determine which residues are likely to be located at the binding interface within a protein complex. Surface residues can participate in molecular interactions with nearby solvent molecules. If these residues are located at the binding interface when part of a complex, upon binding, the side chains of these residues become buried and only interact with the neighboring residues of an adjacent bound subunit, decreasing the number of solvent interactions and the probability of that residue being labeled. In this case, one would expect to observe large changes in the frequency of modification for interface residues between the unbound and bound states of complexes due to large changes in solvent accessibilities in these regions. Based on this hypothesis, large decreases in modification of residues from the unbound to bound state of a complex have a higher probability of participating in the interface.

To test the proposed hypotheses, we used a benchmark set of differential labeling data from three different protein complexes consisting of actin bound to gelsolin segment 1 (actin/gs1, heterodimer, PDB ID: 1YAG),^49^ β-2-microglobulin (homodimer, PDB ID: 2F8O),^50^ and insulin (hexamer of heterodimers, PDB ID: 4INS).^51^ To establish the validity of the hypothesized relationship with experimental data, the native crystal structures of these complexes were used to analyze the proximity of residues to the interface as a function of modification rate in the monomer compared to the complex. We assumed large-scale conformational changes do not occur upon complex formation after examining all subunits of each complex which contained labeled residues and finding that the average RMSD of the unbound to the bound subunits was 2.1 Å. To quantify the amount of change in modification occurring between the unbound/bound state, a modification change was calculated (see methods for full detail) with a larger value indicating a larger decrease in modification from the unbound to bound state. To isolate residues with large modification changes, only residues that saw at least a 40% change in modification between unbound/bound states were considered. Residues that were within 10 Å of the other chain were considered a part of the binding interface. We first used this criterion to compare the native structures to experimental data. The table in Figure 1b lists the number of labeled residues at and outside the interface and those residues are visualized for β-2-microglobulin in Figure 1a. For all three complexes, the majority of labeled residues with modification changes greater than or equal to 40% were found to be located at the binding interface of the complex, with all the designated residues being at the interface for two of the complexes. Overall, 91% of labeled residues with modification changes greater than or equal to 40% were close to protein-protein interfaces. This small preliminary analysis supported our hypotheses and indicated covalent labeling can be used to distinguish interface residues.

Furthermore, we hypothesized that a larger distance to the binding interface for a particular residue would result in less solvent accessibility change when comparing unbound and bound states. For this reason, we would expect a smaller change in modification between the unbound/bound states of a complex. Combining labeling data from all three complexes along with the interface distances of these labeled residues resulted in a more comprehensive analysis as shown in Figure 1c. A linear trend with R^2^ = 0.34 and a normalized root-mean-square error (NRMSE) of 1.5 was observed between modification change (experimental data, y-axis) and the interface distances (native structures, x-axis) for labeled residues. A larger distance between a labeled residue and the other subunits in the bound form correlated with generally smaller changes in modification. This linear correlation was used to predict an expected modification change from any structural model (by calculating the distances to the interface and using the fit line). The linear parameters of slope and intercept from Figure 1c were incorporated within our covalent labeling score term, as described in Methods.

### Structure prediction with covalent labeling data

RosettaDock has had many successes in modeling quaternary protein structure.^55^ And its docking predictive capabilities can be further enhanced with the inclusion of sparse experimental data.^56–58^ The RosettaDock Interface score (Isc) accounts for interactions at the binding interface and can be supplemented with additional score terms to predict more nativelike poses. Here we aimed to explore whether covalent labeling MS data can meaningfully improve model quality. Due do the accuracy of AlphaFold2 (AF2) for monomer prediction, models generated by AF2 were used to provide the input to RosettaDock and a covalent labeling-based score term was used to rescore the oligomeric structures of modeled protein complexes and predict the native structure.

The parameters obtained from the correlation shown in Figure 1c were used to simulate predicted modification changes of labeled residues. For each labeled residue in a modeled complex, the difference between experimental and predicted modification change was calculated and input into a sigmoidal penalty term which penalizes residues with larger disagreement with experimental data (see Equation 3 in Methods). The scores were then summed up for each labeled residue in a model and normalized across all models of a set. The covalent labeling score term was then combined with the Isc to form the covalent labeling score. Since traditional docking consists of two docking partners, the insulin complex was broken up into three separate sub-complexes to model the assembly of all unique interfaces, where AB_CD, ABCD_EFGH, ABCDEFGH_IJKL define what chains made up each docking partner, separated by an underscore. In a first study, we redocked the native crystal structures and Rosetta yielded accurate predictions for 4/5 complexes (Figure S2a). The only exception was β-2-microglobulin, for which a top scoring model with an RMSD of 9.2 Å was identified. When including covalent labeling data in the score function, 5/5 complexes had accurate predictions and the top scoring model for β-2-microglobulin had an RMSD of 3.0 Å (Figure S2b). While these data were promising, the preliminary docking study required crystallographic information of subunit structure in the complex state.

To simulate a more realistic situation, we then used AF2 to generate components (subunits or sub-complexes) of the complexes, which were then input into docking simulations. The top-ranked AF2 models were all accurate with respect to the native structure with RMSD values of 1.4 Å and 0.6 Å for actin and gs1 A and G chains respectively, 1.6 Å for β-2-microglobulin chains, and 1.5 Å for insulin heterodimer. Scoring of the docked sets using covalent labeling data was performed by combining the covalent labeling score term produced by our method with Isc, as previously described. The score versus RMSD plots without using covalent labeling data are shown in Figure 2a, where the top scoring model RMSD with respect to the native structure was 11.2 Å for actin/gs1, 10.1 Å for β-2-microglobulin, 1.7 Å for insulin AB_CD, 9.6 Å for insulin ABCD_EFGH, and 6.8 Å for insulin ABCDEFGH_IJKL. Only 1/5 of the sets of docked structures had a top scoring model with RMSD less than 5 Å. Figure 2b shows the results of docked sets from Figure 2a using our covalent labeling score instead of Isc. Using our score, 5/5 of the sets had top scoring models with an RMSD below 3.6 Å. The top scoring model RMSD with respect to the native structure was 1.6 Å for actin/gs1, 3.17 Å for β-2-microglobulin, 1.73 Å for insulin AB_CD, 3.53 Å for insulin ABCD_EFGH, and 3.54 Å for insulin ABCDEFGH_IJKL. Figure 2c shows the top scoring models for each docked set with the inclusion of our covalent labeling score term aligned to the native crystal structure.

**Figure 2:**
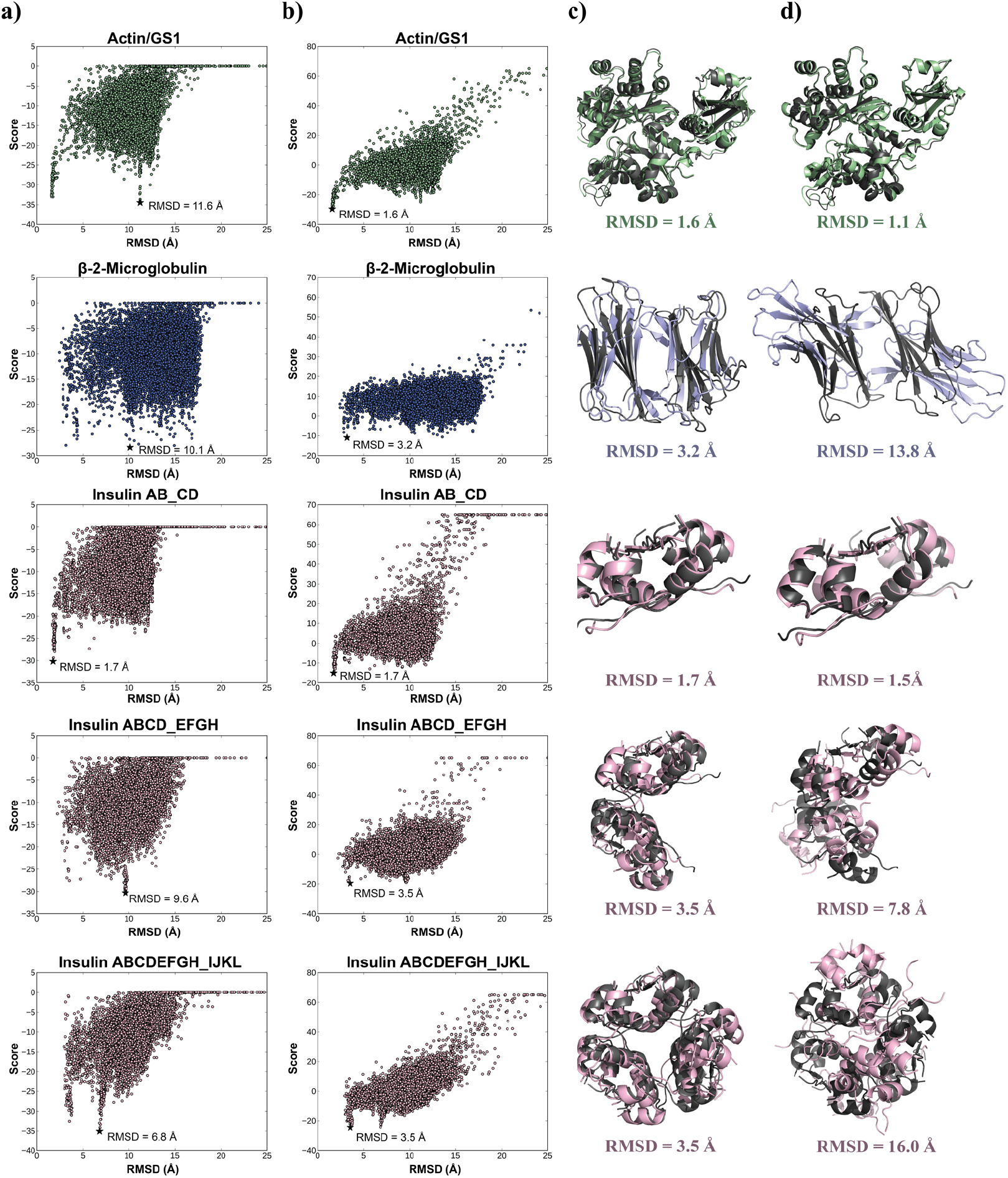
Score vs. RMSD to the crystal structure (a and b) for 10,000 docked models generated for each complex in the benchmark set. The top scoring models are marked by a black star and aligned to crystal structure. Actin/gs1 is shown in green, β-2-microglobulin in blue, and insulin structures in pink. The RMSD of the top scoring model is indicated next to the marked point. a) RosettaDock Isc versus RMSD (without CL data). b) Covalent labeling score versus RMSD (with CL data). c) Top scoring complex model predictions using covalent labeling score aligned to the native crystal structures (dark gray). d) Top ranked models generated by AlphaFold-Multimer aligned to native crystal structure. RMSDs are listed for each complex.

The assessment of additional metrics further demonstrated the benefits of including covalent labeling in scoring. As shown in Table 1, improvements were observed in TM-score and DockQ score upon addition of CL data. TM-score analyzes the topological similarity between structures and DockQ is a quality measure used for evaluation of protein-protein docking data. The average TM-score improved from 0.70 to 0.84 and the average DockQ score improved from 0.21 to 0.50 when including covalent labeling data in scoring. The TM-score and DockQ score for all top scoring models either stayed the same or improved with the addition of experimental data (Table S1). These results demonstrated that the information contained in the covalent labeling modification of residues can indeed facilitate the discrimination of nativelike and non-nativelike poses.

**Table 1:** Average metric analysis for the top scoring models with and without covalent labeling data

|  | Avg RMSD (Å) | Avg TM-Score | Avg DockQ Score |
| --- | --- | --- | --- |
| w/o CL data | 7.89 | 0.70 | 0.21 |
| w/ CL data | 2.70 | 0.84 | 0.50 |

As a comparison to state-of-the-art methodology, we also used AlphaFold-Multimer to predict the full structure of the complexes from our benchmark set without including the native structure as a homolog. Figure 2d shows the generated AlphaFold-Multimer models aligned to the native structures for the complexes. The root mean-squared deviation (RMSD) of the top ranked models for each of the complexes were 1.1 Å, 13.8 Å, 1.5 Å, 7.8 Å, and 16.0 Å for actin/gs1, β-2-microglobulin, insulin AB_CD, insulin ABCD_EFGH, and insulin ABCDEFGH_IJKL, respectively. Only 2/5 complexes in the benchmark set were accurately predicted with an RMSD below 7 Å. Interestingly, for the β-2-microglobulin homodimer, AlphaFold-Multimer predicted accurate individual chains in its top ranked model (with an RMSD of 1.6 Å for both chains) but failed to accurately predict the full complex. It can be seen in Figure 2d that the interface and orientations of the separate chains did not match that of the native structure.

## Conclusion

Sparse experimental data can booster the effectiveness of existing computational techniques. In this current study, we have proposed a hybrid technique utilizing the combination of state-of-the-art computational methods (AlphaFold and RosettaDock) with covalent labeling mass spectrometry data to address cases when the computational tools fail to model accurate complexes. Covalent labeling reagents modify residues based on features such as solvent accessibility, and we have demonstrated that changes in modification of residues in covalent labeling experiments can be used to determine the likely proximity of these residues to the binding interface within protein complexes (Figure 1). As the modification change of a labeled residue between the unbound/bound states of a complex increases, it is more likely to be located at the binding interface. The relationship between experimental modification change and inter-subunit distance was used to predict modification changes of modeled residues. We demonstrated that RosettaDock with the inclusion of our covalent labeling score term can predict accurate models for all the complexes in our benchmark set using AF unbound structures as input. Large improvements in model quality were observed when our score term was included. For example, the RMSD of the top scoring model improved from 11.2 Å to 1.6 Å for actin/gs1 and 10.1 Å to 3.2 Å for β-2-microglobulin (Figure 2a and 2b). This demonstrated that the information contained in the experimental covalent labeling values can improve scoring and model selection within RosettaDock. This score term can be used through the newly developed cl_complex_rescore application within Rosetta. A tutorial for using this application can be found within the Supporting Information. Future work will include increasing the number, valency, and structural types of labeled proteins, along with the types of covalent labeling reagents used, to more comprehensively test the ability of covalent labeling data to elucidate protein complex structure. In this study, we exclusively used differential covalent labeling data since it provides the most accurate structural information. However, many labeling experiments only yield non-differential datasets. In future work, we will focus on developing computational tools that utilize these datasets for complex prediction. In addition, we plan to explore combining other types of complementary experimental MS data with covalent labeling data.

## Acknowledgments

We thank the members of the Lindert lab for many useful discussions and the Ohio Supercomputer Center for valuable computational resources.^59^ Integrative protein modeling work was supported by NIH (P41 GM128577) and a Sloan Research Fellowship to S.L.

