## Supplementary Information for "Protein complex prediction using Rosetta, AlphaFold, and mass spectrometry covalent labeling"

### Supporting Information

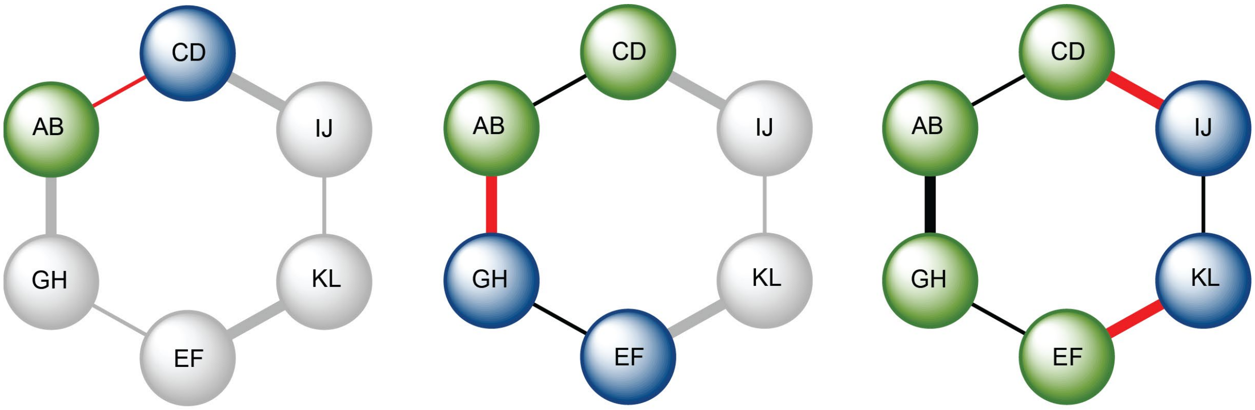


Insulin AB_CD　　Insulin ABCD_EFGH　　Insulin ABCDEFGH_IJKL

**Figure S1:** Simplified diagram illustrating different insulin structures used in our docking benchmark. In different stages, different interfaces were assembled during the docking. Each heterodimer subcomplex is shown as a sphere (with chains labeled). Green and blue spheres indicate the docking partner(s) for each subcomplex. Grey spheres specify which subcomplexes were not included in a particular docking study. Red and black lines indicate predicted and existing interfaces, respectively.

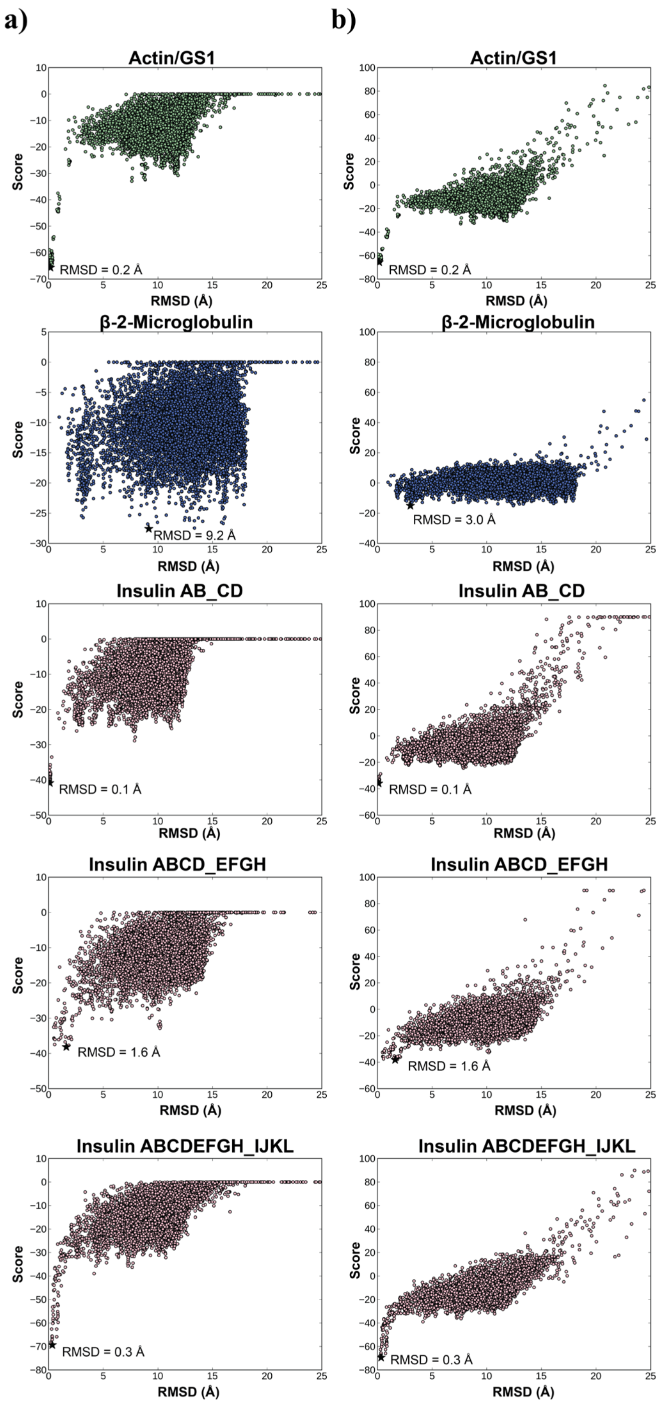


**Figure S2:** Score vs. RMSD to the crystal structure for 10,000 docked models generated for each complex in benchmark set with top-scoring model marked by a black star. Crystal structures were used as docking input. Actin/gs1 is shown in green, β-2-microglobulin in blue, and insulin structures in pink. The RMSD of the top-scoring model is indicated next to the marked point. a) RosettaDock Isc versus RMSD (without CL data). b) Covalent labeling score versus RMSD (with CL data).

**Table S1:** Model quality metric analysis for the top scoring models of each docked set with and without covalent labeling data.

| Metric | Complex | w/o CL data | w CL data |
|---|---|---|---|
| RMSD (Å) | Actin/GS1 | 11.20 | 1.60 |
| | Beta | 10.10 | 3.20 |
| | Insulin AB_CD | 1.73 | 1.70 |
| | Insulin ABCD_EFGH | 9.60 | 3.50 |
| | Insulin ABCDEFGH_IJKL | 6.80 | 3.50 |
| TM-Score | Actin/GS1 | 0.78 | 0.97 |
| | Beta | 0.53 | 0.75 |
| | Insulin AB_CD | 0.89 | 0.89 |
| | Insulin ABCD_EFGH | 0.55 | 0.74 |
| | Insulin ABCDEFGH_IJKL | 0.74 | 0.83 |
| DockQ Score | Actin/GS1 | 0.07 | 0.65 |
| | Beta | 0.10 | 0.44 |
| | Insulin AB_CD | 0.70 | 0.70 |
| | Insulin ABCD_EFGH | 0.06 | 0.36 |
| | Insulin ABCDEFGH_IJKL | 0.12 | 0.34 |

**cl_complex_rescore Rosetta Application Tutorial**

This application is designed to rescore large sets of docked models with differential covalent labeling data to improve the quality of the top-scoring model. The application can be used for situations such as a generating a structure for a protein complex of interest which has been studied using covalent labeling but does not have an existing crystal structure. This tutorial consists of two parts: generating docked models using RosettaDock and rescoring those models with the cl_complex_rescore application.

Subunits for a particular complex of interest can be generated using AlphaFold2 (as we did in this manuscript). AlphaFold2 can be installed by following the instructions available at https://github.com/deepmind/alphafold. Once installed, the structure of a subunit can be built by providing AlphaFold2 the protein sequence as a FASTA file. If the subunit is comprised of multiple chains, the `--model_preset` option should be changed to multimer (the default is monomer). However, this is only one approach to obtaining structures of the subunits. Subunits can be built using approaches other than AlphaFold2 such as *ab initio* modeling or homology modeling.

In this tutorial, variables that need to be specified by the user are shown in brackets (< >):

**Part 1: Protein-Protein Docking with RosettaDock**

1. Prepare a pdb file containing generated subunit models.
2. Prepack the chains by running the following command:
   ```
   ~/Rosetta/main/source/bin/docking_prepack_protocol.default.<os><compiler>release -in:file:s <pdb> -partners <chains>
   ```
   - <os> operating system (macOS, linux).
   - <compiler> compiler used (gcc, clang, etc.).
   - <pdb> path and filename of pdb file created in step 1.
   - <chains> chains of the docking partners, separated by underscore. Ex: A_B where A is the static chain, B is the mobile chain.

   Example: Using linux OS, gcc as compiler, model.pdb as pdb file, and docking chains A and B.

   ```
   ~/Rosetta/main/source/bin/docking_prepack_protocol.default.linuxgccrelease -in:file:s model.pdb -partners A_B
   ```
3. To dock the chains using RosettaDock, use the following command:
   ```
   ~/Rosetta/main/source/bin/docking_protocol.default.<os><compiler>release -in:file:s <prepacked_pdb> -partners <chains> -nstruct <n_structs>
   ```
   - <prepacked_pdb> the output pdb from previous step
   - <n_structs> The number of models to be generated (≥10,000 recommended)
   - Additional flags can be specified, such as a perturbation flag (randomize1, -randomize2, -spin, etc.) which can be used to search more conformational space. A list of all available flags and their relevant details are available at

https://www.rosettacommons.org/docs/latest/application_documentation/docking/docking-protocol

Example: Dock 10,000 structures of chains A and B of ppk_model_0001.pdb

```
~/Rosetta/main/source/bin/docking_protocol.default.linuxgccrelease -in:file:s ppk_model_001.pdb -partners A_B -nstruct 10000
```

**Part 2: cl_complex_rescore**

1. Create a text file listing the paths to each of the models/structures you wish to rescore with the application.
   Example:
```
/path/to/file/dock_ppk_model_0001_00001.pdb
/path/to/file/dock_ppk_model_0001_00002.pdb
/path/to/file/dock_ppk_model_0001_00003.pdb
/path/to/file/dock_ppk_model_0001_00004.pdb
/path/to/file/dock_ppk_model_0001_00005.pdb
…
/path/to/file/dock_ppk_model_0001_10000.pdb
```
2. Create a second text file listing each labeled residue and the degree of modification for each labeled residue for both the unbound and bound states as well as which chain(s) the residue is located on. The first column should indicate the sequence number of each residue (Note: number residues by treating the first residue of each chain as being at position 1). The second and third column should contain the modification rate (or other metric of modification) of the unbound and bound state of the complex, respectively. The fourth column should contain each chain the residue is located on (no spaces).
   Example*:

```
#Residue Index   Unbound Modif.   Bound Modif.       Chains
7                56.4             34.7               ACEGIK
8                56.4             34.7               ACEGIK
10               56.4             34.7               ACEGIK
13               56.4             34.7               ACEGIK
14               56.4             34.7               ACEGIK
19               56.4             34.7               ACEGIK
1                1.4              0.4                BDFHJL
5                14.9             7.6                BDFHJL
7                14.9             7.6                BDFHJL
10               14.9             7.6                BDFHJL
16               8.6              1.8                BDFHJL
17               8.6              1.8                BDFHJL
22               8.6              1.8                BDFHJL
24               9.3              5.1                BDFHJL
25               9.3              5.1                BDFHJL
26               9.3              5.1                BDFHJL
```

| | | | |
|---|---|---|---|
| 28 | 0.55 | 0.23 | BDFHJL |

* The '#' header row is not required.

3. Once the necessary text files have been generated, the models can be rescored using the cl_complex_rescore application by running the following command.
```
~/Rosetta/main/source/bin/cl_complex_rescore.default.<os><compiler>release -in:file:l <file_with_paths_to_models> -cl_data <data_file> -interface <chains>
```
   - <file_with_paths_to_models> path and filename of list of files created in step 1.
   - <data_file> path and filename of file created in step 2.
   - <chains> chains defining the interface, separated by underscore. Ex: A_B or AB_C.

   Example: Using linux OS, gcc as compiler, pdblist.txt (list of pdb paths), insulin_data.txt (CL data file) and the interface is between chains ABCDEFGH and IJKL.
```
~/Rosetta/main/source/bin/cl_complex_rescore.default.linuxgccrelease -in:file:l pdblist.txt -cl_file insulin_data.txt -interface ABCDEFGH_IJKL
```

After successfully running the application, a "cl_complex_score.out" file will be generated. The first column specifies the model, the second column is the raw penalty score for the model (before normalization and unweighted), and the third column is the weighted and normalized penalty score.

Example output file:
```
Model                          Raw_Penalty    Weighted_CL_Score_Term
input_files/model_00001.pdb    7.71483        12.2308
input_files/model_00002.pdb    7.48138        11.8607
...
input_files/model_10000.pdb    7.00656        11.108
```
